## Supporting Information for "Fluorescence-based mapping of condensate dielectric permittivity uncovers hydrophobicity-driven membrane interactions"

*Supplementary Information*

E. Sabri[1], A. Mangiarotti[1,2,3] and R. Dimova[1]

[1]*Max Planck Institute of Colloids and Interfaces, Science Park Golm, 14476 Potsdam, Germany*
[2]*Centro de Investigaciones en Química Biológica de Córdoba (CIQUIBIC), CONICET, X5000HUA Córdoba, Argentina.*
[3]*Departamento de Química Biológica Ranwel Caputto, Facultad de Ciencias Químicas, Universidad Nacional de Córdoba, X5000HUA Córdoba, Argentina.*

### Derivation of the adapted Lippert-Mataga formula

The derivation of Eq. 1 in the main text is based on fluorescence first-principles detailed in ref. [1]. Briefly, it considers a point dipole (the fluorophore) embedded in a continuous dielectric medium (solvent), where the red shift in fluorophore emission arises from dipole-dipole coupling between the fluorophore and the solvent molecules. This interaction is governed by the energy of a dipole molecule in an external field given by

$$E_{dipole} = \mu_d R, \qquad \text{(S1)}$$

where $\mu_d$ and $R$ are the dipole moment and the external electric field. Additionally, the reaction electric fields of the solvent with respect to the fluorophore ground and excited state dipoles are given by

$$R_{el}^{G} = \frac{\mu_G}{V_{cav}} f(n), \qquad \text{(S2)}$$

$$R_{el}^{E} = \frac{\mu_E}{V_{cav}} f(n), \qquad \text{(S3)}$$

$$R_{dip}^{G} = \frac{\mu_G}{V_{cav}} (f(\varepsilon) - f(n)), \qquad \text{(S4)}$$

$$R_{dip}^{E} = \frac{\mu_E}{V_{cav}} (f(\varepsilon) - f(n)), \qquad \text{(S5)}$$

where $R_{dip}^{G}$, $R_{dip}^{E}$, $R_{el}^{G}$, $R_{el}^{E}$, $\mu_G$, $\mu_E$, ε, $n$, $V_{cav}$ and $f$ respectively represent the reaction fields associated with solvent dipolar relaxation near the ground and excited dipole fluorophore, the reaction fields associated with the electronic rearrangement of the solvent molecules around the ground and excited state dipoles, the ground-state dye dipole, the excited-state dye dipole, solvent permittivity, solvent refractive index, the volume of the virtual spheroidal cavity in which the dye molecule resides and the polarizability function that depends on the geometry of the cavity [1, 2].

The ground- and excited-state energies associated with photon absorption by the fluorophore (see Fig.1a) read

$$E_{abs}^{G} = (E_{abs}^{G})_v - \mu_G R_{dip}^{G} - \mu_G R_{el}^{G}\ , \qquad \text{(S6)}$$

$$E_{abs}^{E} = (E_{abs}^{E})_v - \mu_E R_{dip}^{G} - \mu_E R_{el}^{E}, \qquad \text{(S7)}$$

where $(E_{abs}^{G})_v$ and $(E_{abs}^{E})_v$ respectively correspond to the absorption energy levels of the ground and excited state fluorophore in vacuum.

The ground- and excited-state energies associated with photon emission by the fluorophore (see Fig.1a) are

$$E_{em}^{G} = (E_{em}^{G})_v - \mu_G R_{dip}^{E} - \mu_G R_{el}^{G}\ , \qquad \text{(S8)}$$

$$E_{em}^{E} = (E_{em}^{E})_v - \mu_E R_{dip}^{E} - \mu_E R_{el}^{E}, \qquad \text{(S9)}$$

where $(E^G_{em})_v$ and $(E^E_{em})_v$ respectively correspond to the emission energy levels of the ground and excited states of the dye molecule in vacuum.

Because the energy of the absorbed and emitted photons corresponds to the gap between the ground and excited energy levels, the wavenumbers of absorbed and emitted wave trains read

$$\nu_a = \frac{E^E_{abs}-E^G_{abs}}{hc} = \mu_G \frac{\mu_G-\mu_E}{V_{cav}hc}\left(f(\varepsilon)-f(n)\right) + \frac{\mu_G^2-\mu_E^2}{V_{cav}hc} f(n) + \frac{(E^E_{abs})_v-(E^G_{abs})_v}{hc}, \qquad \text{(S10)}$$

$$\nu_f = \frac{E^E_{em}-E^G_{em}}{hc} = \mu_E \frac{\mu_G-\mu_E}{V_{cav}hc}\left(f(\varepsilon)-f(n)\right) + \frac{\mu_G^2-\mu_E^2}{V_{cav}hc} f(n) + \frac{(E^E_{em})_v-(E^G_{em})_v}{hc}, \qquad \text{(S11)}$$

where $h$ and $c$ respectively are the Planck constant and the speed of light in vacuum. By setting $(\nu_f)_v = \frac{(E^E_{em})_v-(E^G_{em})_v}{hc}$ and assuming $f(n^2) \sim f(n_{H_2O}{}^2)$ for all considered solvents we obtain

$$\nu_f = \mu_E \frac{(\mu_G-\mu_E)}{V_{cav}hc} f(\varepsilon) + \left[(\nu_f)_v + f(n_{H_2O}{}^2)\frac{\mu_G(\mu_G-\mu_E)}{V_{cav}hc}\right], \qquad \text{(S12)}$$

where the second term is a constant and $\nu_f$ is a single variable function of $\epsilon$. By setting $\left[(\nu_f)_v + f(n_{H_2O}{}^2)\frac{\mu_G(\mu_G-\mu_E)}{V_{cav}hc}\right] = \text{const}$, we recover Eq. 1 in the main text.

The generalized Debye function $f$ has the following expression [2, 3]

$$f(x) = \frac{\beta(1-\beta)(x-1)}{x+(1-x)\beta}, \qquad \text{(S13)}$$

where $\beta$ is a constant that depends on the length of the three principal symmetry axes of the spheroidal molecule. As such, Eq. S12 can be rewritten as

$$\nu_f = \alpha \frac{\beta(1-\beta)(\epsilon-1)}{\epsilon+(1-\epsilon)\beta} + \gamma, \qquad \text{(S14)}$$

where $\alpha, \beta$ and $\gamma$ are three fitting parameters. Here, to avoid making further *ad hoc* assumptions beyond that of the spheroidal shape of the ACDAN dye molecule, we use Eq. S14 to fit the data presented in Fig. 1(c-d). All fitting parameters and relevant physical quantities are provided in Table S2. Additionally, the relation $\frac{\phi(\lambda_f-\lambda_0)}{2\pi} + \lambda_0 = \lambda$ was used to establish the link between the phase $\phi$ and the emission wavelength $\lambda$ for the data presented in Fig. 1d and S1f.

**Spectrofluorimetry of ACDAN for solvent permittivity measurements and comparison of sensitivity to hyperspectral imaging**

Bulk fluorescence measurements were made using a FluoroMax4® (HORIBA Scientific) fluorimeter with the FluorEssence (V3.5) software. The excitation wavelength was set to 360nm with an excitation slit width of 5nm and a detection slit width of 2nm. Data acquisition was performed at 25°C using a 1nm detection wavelength increment over a (375-700) nm spectral window for mineral oil samples and over a (400-700) nm spectral window for the remaining samples. Quartz chambers (Hellma® Analytics) were used to perform all measurements, chamber volumes of 50µL were used for measuring the emission of ACDAN in water-dioxane mixtures, ethanol 100% and butanol 100%, 200µL for measuring the emission of ACDAN in PEG400-water mixtures and 2.8mL for measuring the emission of ACDAN in anisol-hexanol mixtures and mineral oil. The measurement of each condition consisted in three independent trials which were then averaged and the error was computed by performing an error propagation analysis accounting for the slit parameters and the standard deviations associated with each trial triplet. The ACDAN concentration was 1µM for all measurements.

***Spectrofluorimetry analysis***

While spectrofluorimetry offers high spectral resolution, it captures the bulk fluorescence response of a solution, precluding spatial differentiation. Consequently, it cannot distinguish between coexisting phases in heterogeneous samples, thereby limiting its ability to resolve the distinct dielectric properties of individual components. However, its spectral sensitivity is often higher than conventional microscopy detectors[3]. Figure S1(a-c) shows the fluorescence emission of ACDAN in various solvents and homogeneous mixtures with permittivities ranging from $2\varepsilon_0$ to $80\varepsilon_0$.

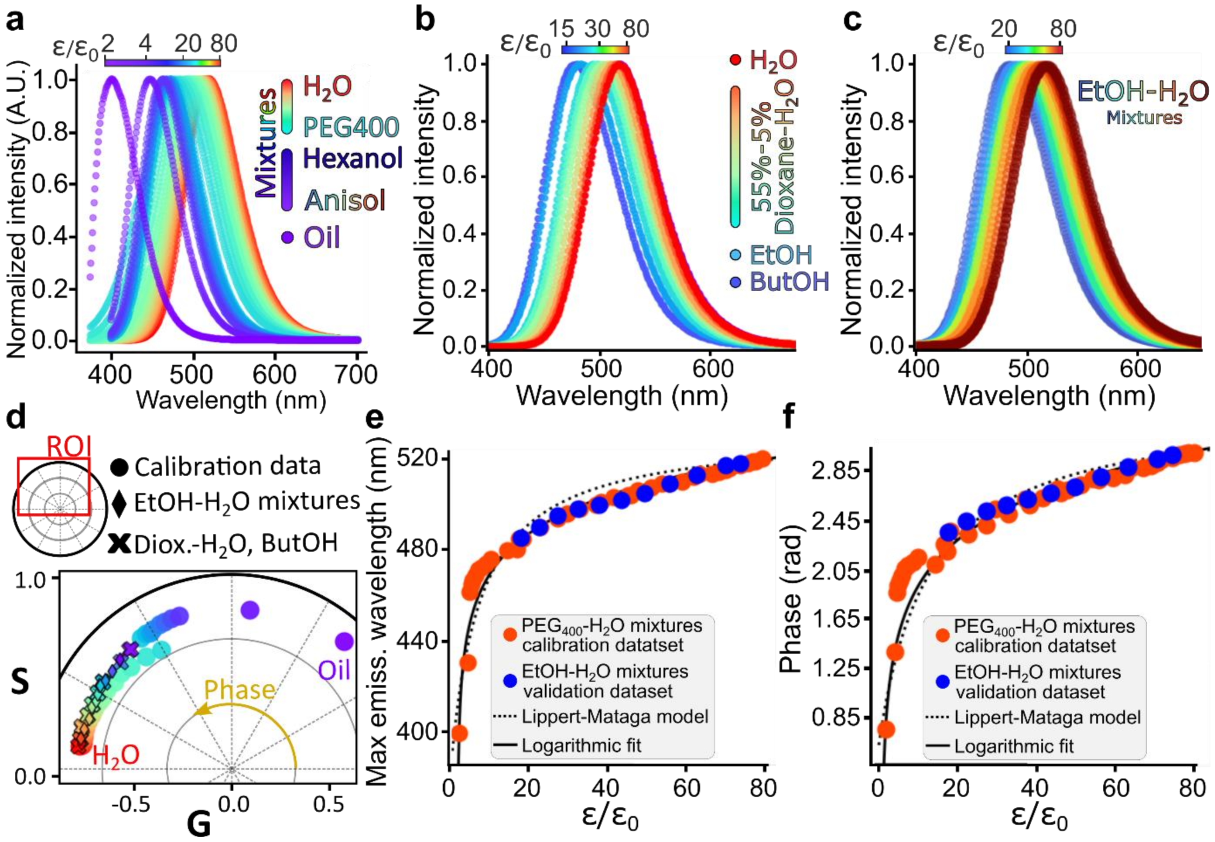


**Figure S1: ACDAN fluorescence spectroscopy enables high-accuracy and broad-range quantitative permittivity measurements. (a)** Spectrofluorimetry measurements of ACDAN fluorescence in homogeneous solvents including mineral oil, various mixtures of anisol and 1-hexanol, and different PEG-400 solutions in water (the mixture and solution concentrations are given in Table S1). The color bar above the graph represents the dielectric constants of these mixtures taken from the literature[4-6] also given in Table S1. **(b)** Emission spectra of ACDAN in organic solvents of different permittivities. **(c)** Emission spectra of ACDAN in ethanol-water mixtures of 5%, 10%, 20%, 30%, 40%, 50%, 60%, 70%, 80%, 90% and 100% ethanol content. **(d)** Phasor plot signatures corresponding to the emission spectra of the calibration solutions presented in (a) (dots) as well as the solvents presented in (a-b) (crosses and diamonds respectively). The bounds ($\lambda_0$; $\lambda_f$) of the phasor plot were set to (370; 700) nm. **(e-f)** Calibration data (orange circles) showing (e) the maximum emission wavelength and (f) the average phase value for ACDAN in the solutions introduced in panels (a-c) (dioxane-water mixtures, ethanol and butanol) and to the permittivity for each solution, plotted as a function of dielectric constants reported in the literature[5, 7] and referenced in Table S1. Each data point represents the mean±SD of three measurements (n=3); SD values are smaller than the symbol size. The dotted line represents the Lippert-Mataga equation (Eq. 1 in the main text) and the solid curve shows a logarithmic fit (Eq. 2) used as a calibration curve. Additional data for ethanol-water mixtures (blue circles) are included, where the dielectric constants were calculated using the Maxwell-Garnett law of mixtures (see Table S3 and panel (c)). In (f), the relation $\frac{\phi(\lambda_f-\lambda_0)}{2\pi}+\lambda_0=\lambda$ with $\lambda_0=375$nm and $\lambda_f=700$nm was used to establish the link between the phase $\phi$ and the emission wavelength $\lambda$.

Figure S1d demonstrates qualitative agreement between the Lippert-Mataga equation (Eq. 1 in the main text) and our experimental data across a broad range of systems: mineral oil, ethanol, butanol, various mixtures of anisol and 1-hexanol, dioxane-water, and different PEG-400 solutions in water. The small discrepancy between the experimental data and the theoretical description provided by Eq. 1 likely stems from the model's simplified assumptions regarding ACDAN's molecular geometry[1, 2]; specifically, the spherical shape assumed in the model does not account for the rotational degrees of freedom at the C-N and C-C bonds of ACDAN's extremities. Additionally, the ground and excited states of the dye dipole are assumed to be constants, independent of the solvent molecules surrounding the dye. This assumption implicitly suggests that the electronic and dipolar reaction fields associated with solvent molecules are the only electrochemical conformational rearrangements that can occur in the dye-solvent system. However, numerical simulations presented in refs. [3, 8], based on Monte-Carlo sequential quantum mechanics and molecular mechanics approaches[9], crossed compared with the polarizable continuum model[10], suggest that the ground state dipole of PRODAN (differing from the ACDAN only by a $CH_2$ at its ethanal extremity) can increase by up to 40% between vacuum and aqueous environments. We believe that these caveats in the assumptions of the Lippert-Mataga framework explain the observed discrepancy between our measurements its prediction in Fig. S1d.

To further refine the calibration, we proposed a logarithmic fit function (legends of Fig. S1d and f), which improves the empirical agreement with spectrofluorimetry datasets (Fig. S1d and f).

#### *Hyperspectral imaging analysis*

Figure S2 presents the calibration approach used to compute the dielectric permittivity of different samples for all microscopy-based hyperspectral imaging measurements performed in this study. Figs. S2a-b show respectively a phasor representation and the associated phase histograms of the pixel clouds associated with the calibration solutions (see Fig. 1d), and maps of the individual measurements are presented in Fig. S2c.

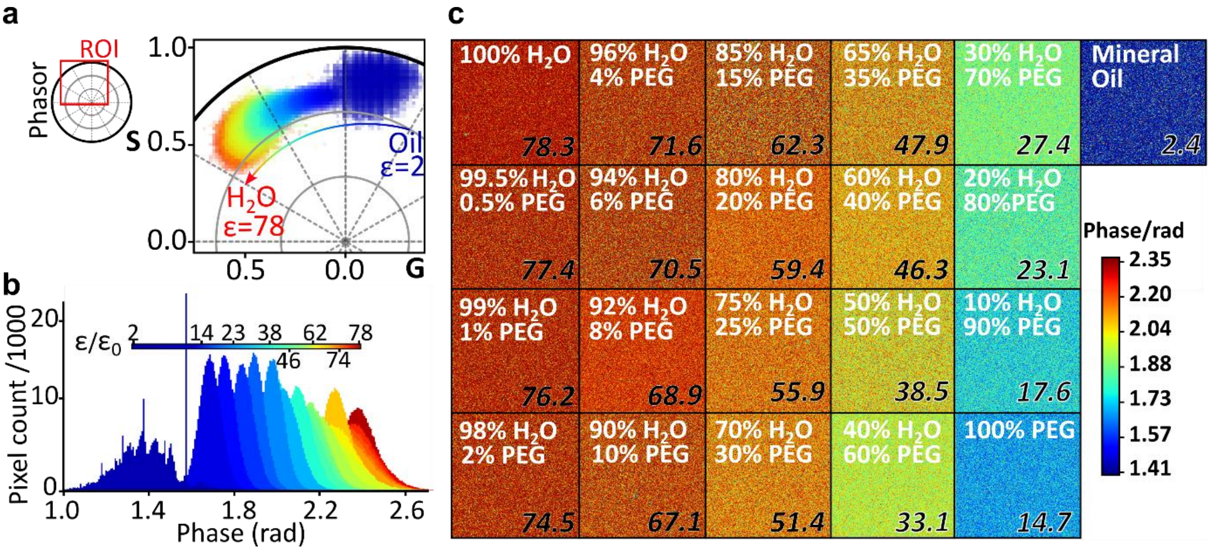


**Figure S2: Calibration procedure of microscopy-based permittivity measurements. (a)** Phasor plot of ACDAN emission in different calibration solutions. **(b)** Phase histograms of the different calibration solutions. The color bar represents the rescaled permittivity of each solution which were taken from the literature. **(c)** Phase map of the different mixtures used as calibration solutions. Each map corresponds to a 36.9x36.9μm$^2$ acquisition frame. Rescaled permittivities are given in the bottom right corner of each frame.

To validate the equivalence of these data analysis approaches as well as the consistency between measurements from different instruments (microscope versus spectrofluorimeter), Fig. S3 provides a comparison between the permittivity values for water-ethanol mixtures obtained using both the DFT and Gaussian fit approaches.

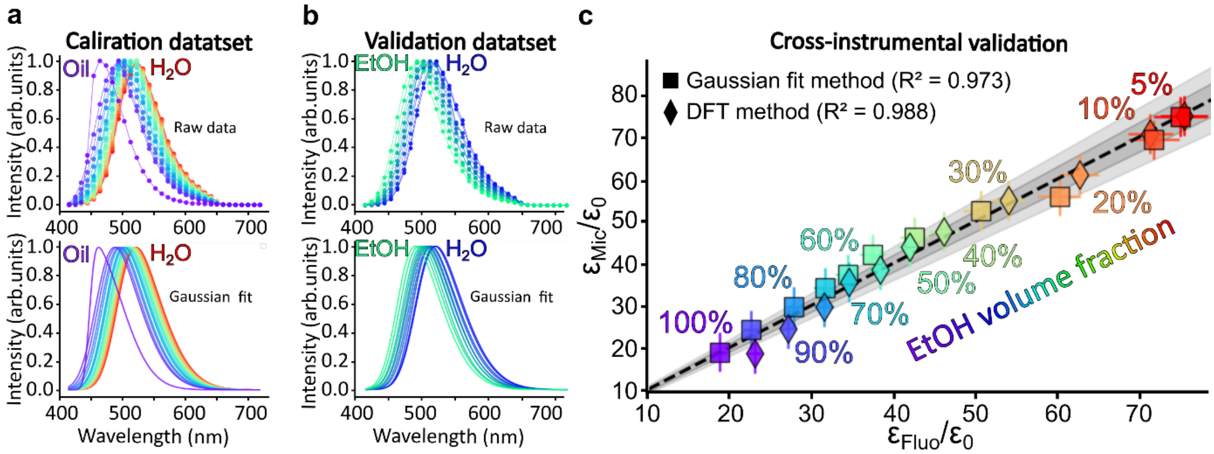


**Figure S3: ACDAN hyperspectral imaging and spectrofluorimetry yield comparable sensitivity to solvent permittivity regardless of the analytical framework. (a,b)** Fluorescence emission profiles of ACDAN in calibration solutions of (b) mineral oil or PEG400-$H_2O$ solutions (100%, 90%, 80%, 70%, 60%, 50%, 45%, 40%, 35%, 30%, 25%, 20%, 15%, 10%, 8%, 6%, 4%, 2%, 1%, 0.5%, 0% PEG content) and (c) ethanol-water mixtures (5%, 10%, 20%, 30%, 40%, 50%, 60%, 70%, 80%, 90% and 100% ethanol content). Top panels: raw data. Bottom panels: Gaussian fits. **(c)** Cross-comparison of permittivity values obtained by spectrofluorimetry (horizontal axis) and hyperspectral microscopy (vertical axis) for the $H_2O$-EtOH mixtures shown in (b). Data points cluster tightly around the identity line (slope = 1), demonstrating excellent agreement between the two experimental methods. Light and dark grey shaded areas indicate 10% and 5% deviation intervals, respectively. The two spectral analysis approaches, DFT (diamonds) and Gaussian fit (squares), yield nearly identical results ($R^2 > 0.97$ for a linear correlation) confirming the robustness and consistency across the methods. Symbols color represents ethanol concentration, corresponding to the respective volume percentages indicated.

Fig. S3(a-b) present the calibration and validation datasets used to perform this cross-comparison. Despite expected differences in calibration curve profiles due to instrument-specific spectral responses [11] (compare Figs. S1d and 1c in the main text), the results presented in Fig. S3c align very well, with data clustering around the identity line (slope 1) and yielding high coefficients of determination ($R^2 > 0.97$). This strong agreement between the two analytical approaches and across different instrumental platforms confirms the instrumental versatility of our calibration approach.
The Gaussian fitting and DFT approaches are presented in the main text and allow to compute the dielectric permittivity based on respectively the position of the maximum wavelength of emission $\lambda_{max}$ (see Fig. 1b-c and additional data and examples in Figs. S1(a-d) and S3(a-b)) and on the phase of the phasor representation of the emission spectrum (see Figs. S1d,f and Fig. 1d).

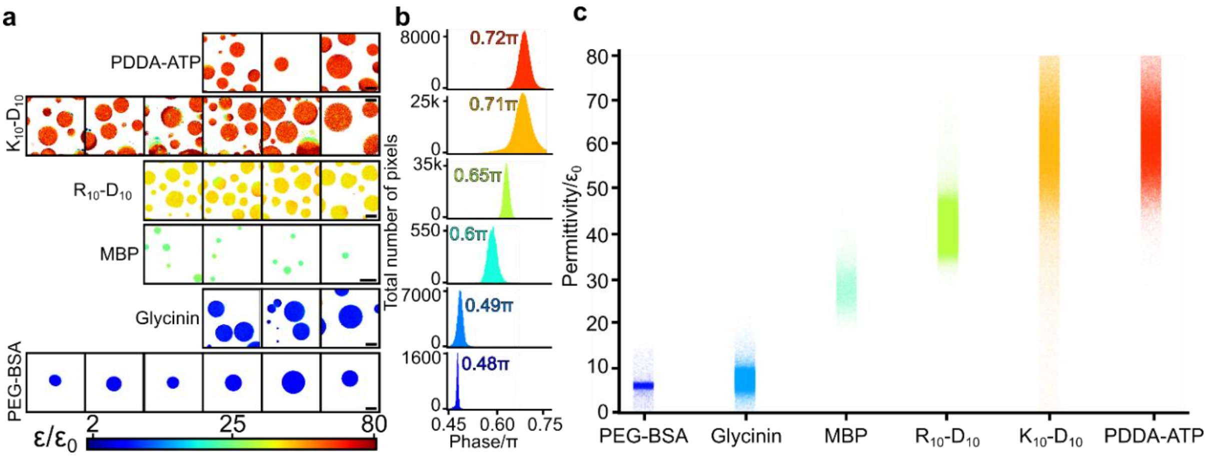


**Figure S4: Pixel-level computation of condensate permittivity**. **(a)** Permittivity mapping of different condensate samples introduced in Fig. 2. Scalebars represent 10 µm. The color bar represents the permittivity on a logarithmic scale. The final concentration of the different compounds composing each solution were of : *PDDA-ATP*, 14.8mM ATP in water, 4.9mM PDDA in water; *$K_{10}$-$D_{10}$*, 2.5mM $K_{10}$ in water, 2.5mM $D_{10}$ in water; *$R_{10}$-$D_{10}$*, 2.5mM $R_{10}$ in water, 2.5mM $D_{10}$ in water; *MBP*, 2.5g/L MBP in water, 10mM NaOH in water; *Glycinin*, 10g/L glycinin in water, 100mM NaCl in water; *PEG-BSA*, 10% PEG-8000 in PBS, 0.5mM BSA in PBS. **(b)** Histograms of the different phases of the total number of pixels represented in the maps of (a). The mean phase values are noted for each distribution. **(c)** Permittivity strip plot of all pixels represented in the histograms of (b). The larger scatter in some of the samples is due to movement (coalescence and sedimentation during acquisition) of condensates lacking strong density contrast with the environment.

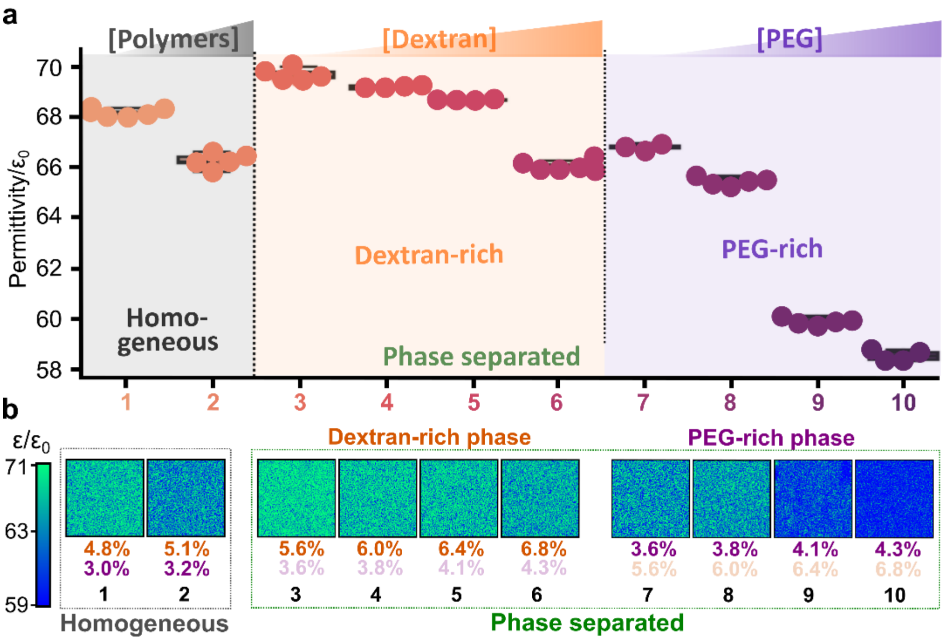


**Figure S5: Permittivity of PEG-dextran aqueous two-phase systems.** (a) Permittivities of homogeneous and phase separated solutions at different total polymer concentration. Black, orange and purple data respectively represent homogeneous, dextran-rich and PEG-rich solutions. Hyperspectral imaging-based measurement of the permittivity of the PEG-dextran mixture compositions in Fig. 3a,b, see Table S4 for exact compositions of the PEG- and dextran-rich phases. (b) Respective permittivity maps and polymer weight fractions: orange and magenta percentages represent the dextran and PEG concentrations respectively. The color bar on the left represents the rescaled permittivity on a logarithmic scale. Each map represents a 36.9x36.9µm$^2$ acquisition frame. The numbers below each condition relates to the referencing of points in the phase diagram in Fig. 3a-b of the main text.

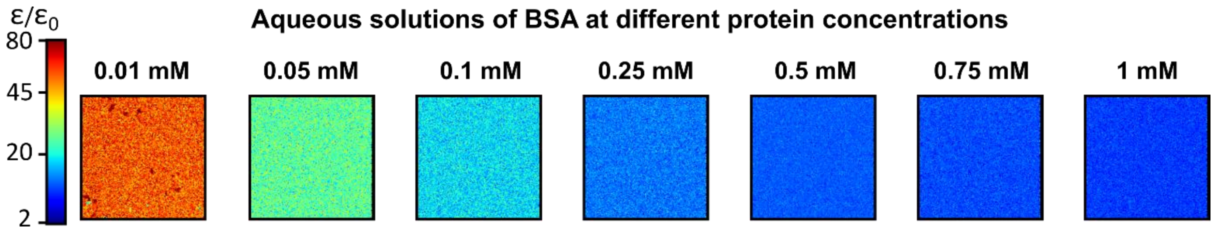


**Figure S6: Permittivity map of BSA solutions of different concentrations.** All vertically aligned frames share the same concentration conditions. The color bar represents the rescaled permittivity. Each map represents a 36.9x36.9μm$^2$ acquisition frame.

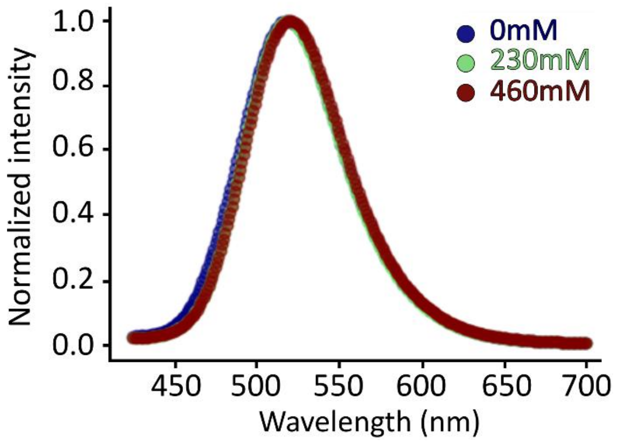


**Figure S7: Control for salt effects on ACDAN-reported permittivity.** Spectrofluorimetric measurements of ACDAN fluorescence emission in glycinin-free aqueous solutions at increasing NaCl concentrations (legend). No systematic spectral changes are observed. The corresponding permittivity values vary by less than ~4$\varepsilon_0$ across the explored salt range (80±2$\varepsilon_0$ for 0 mM, 78±5$\varepsilon_0$ for 230mM and 82±2$\varepsilon_0$ for 460mM NaCl), demonstrating that NaCl alone does not significantly affect the permittivity reported by ACDAN.

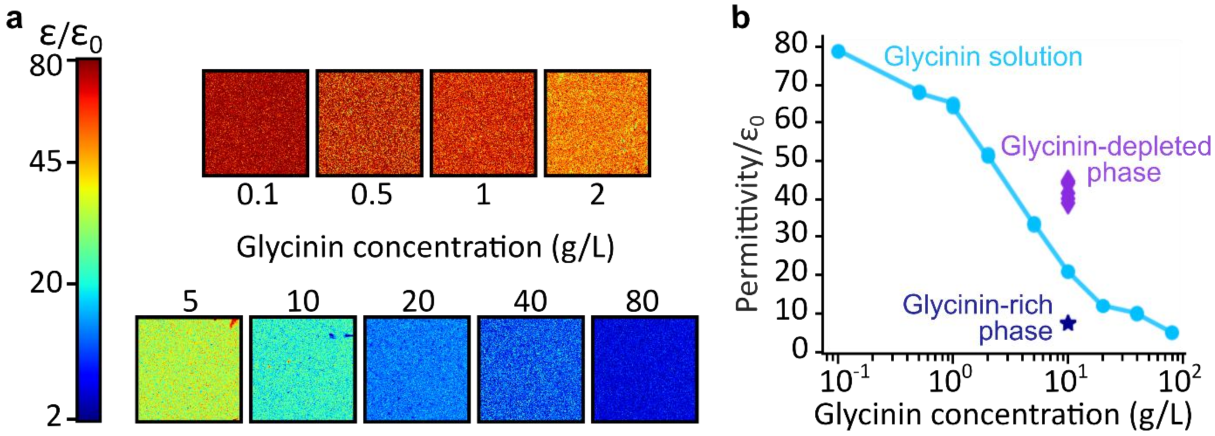


**Figure S8: Comparing the permittivity of glycinin solutions with each phase of a glycinin phase-separated sample. (a)** Mapping the permittivity of glycinin aqueous solutions at different concentrations. Each map represents a 36.9x36.9μm$^2$ acquisition frame. The color bar represents the rescaled permittivity on a logarithmic scale. **(b)** Comparison between the permittivity of glycinin aqueous solution and that of a phase separated sample of 10 g/L glycinin and 100mM NaCl (point 2 of Fig. 3c in the main text). The glycinin-depleted phase at a total glycinin concentration of 10 g/L exhibits a permittivity similar to that of a 3 g/L glycinin solution, suggesting that the latter closely approximates the concentration of the depleted phase. Similarly, the dense phase permittivity suggests that it is approximated by 60 g/L glycinin solution.

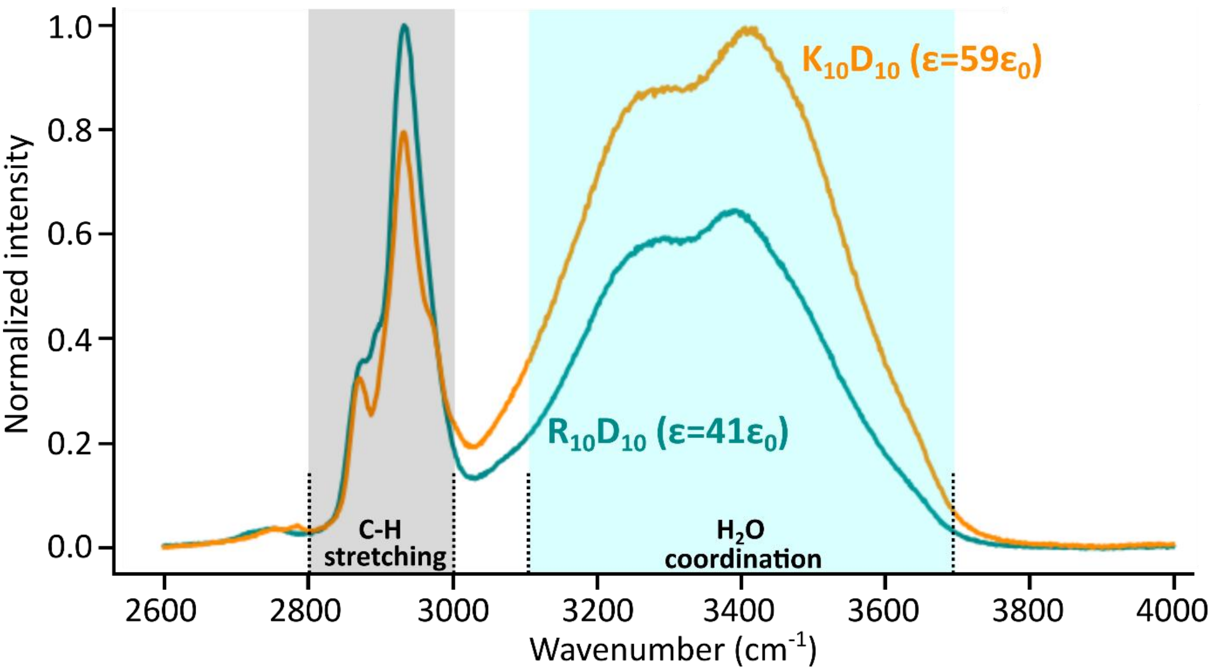


**Figure S9: Raman spectra of peptide-based condensate systems of different permittivity.** The grey region corresponds to the protein C-H bond stretching band (2800-3000 cm$^{-1}$) [12] and the light-blue region corresponds to the water band (3100-3700 cm$^{-1}$) [13]. Compared to $R_{10}$-$D_{10}$ condensates, $K_{10}$-$D_{10}$ condensates contain higher portion of water relative to peptide. The final peptide concentrations were of 2.5mM $D_{10}$ and 2.5mM $K_{10}$ (respectively $R_{10}$) for the $K_{10}$-$D_{10}$ (respectively $R_{10}$-$D_{10}$) condensate solution. This finding aligns with the observed $\sim 20\varepsilon_0$ increase in the permittivity of $K_{10}$-$D_{10}$ condensates shown in Fig. 2 in the main text.

Raman spectra were acquired with a Raman confocal microscope Alpha300 R (WITec GmbH, Germany) with Zeiss EC Epiplan 50×, 0.75NA objective, at an excitation wavelength of 532 nm and 50 mW laser power. Spectra were acquired in the range 400–4100 cm$^{-1}$, each acquisition consisted of an average of seven successive acquisitions performed over a 10s integration timespan. The Raman band of the silicon wafer was used to calibrate the spectrometer. Data were analyzed with the Project FIVE v.5.2 data evaluation software from WITec, and the background was interpolated and subtracted using the OriginPro2023 software via the built-in baseline subtraction procedure.

**Table S1:** Numerical values of different solvents used as calibration data to construct the $\lambda - \varepsilon$ calibration curves used in this study and based on ε measurements available in the literature. All solvent concentrations are in vol%.

| Solvent | $\varepsilon/\varepsilon_0$ | Figures | Reference |
|---|---|---|---|
| $H_2O$ | 78.3 ± 0.5 | 1, S1, S2, S3 | [4] |
| 99.5% $H_2O$ - 0.5% PEG400 | 77.4 ± 0.5 | 1, S1, S2, S3 | [4] |
| 99% $H_2O$ - 1% PEG400 | 76.1 ± 0.5 | 1, S1, S2, S3 | [4] |
| 98% $H_2O$ - 2% PEG400 | 74.5 ± 0.5 | 1, S1, S2, S3 | [4] |
| 5% Dioxane - 95% $H_2O$ | 74 ± NA | 1, S1 | [7] |
| 96% $H_2O$ - 4% PEG400 | 71.6 ± 0.5 | 1, S1, S2, S3 | [4] |
| 94% $H_2O$ - 6% PEG400 | 70.5 ± 0.5 | 1, S1, S2, S3 | [4] |
| 92% $H_2O$ - 8% PEG400 | 68.9 ± 0.5 | 1, S1, S2, S3 | [4] |
| 90% $H_2O$ - 10% PEG400 | 67.1 ± 0.5 | 1, S1, S2, S3 | [4] |
| 15% Dioxane - 85% $H_2O$ | 64 ± NA | 1, S1 | [7] |

| Solvent | ε/ε₀ | Figures | Reference |
|---|---|---|---|
| 85% $H_2O$ - 15% PEG400 | 62.3 ± 0.4 | 1, S1, S2, S3 | [4] |
| 80% $H_2O$ - 20% PEG400 | 59.4 ± 0.4 | 1, S1, S2, S3 | [4] |
| 75% $H_2O$ - 25% PEG400 | 55.9 ± 0.4 | 1, S1, S2, S3 | [4] |
| 25% Dioxane - 75% $H_2O$ | 54 ± NA | 1, S1 | [7] |
| 70% $H_2O$ - 30% PEG400 | 51.4 ± 0.4 | 1, S1, S2, S3 | [4] |
| 35% Dioxane - 65% $H_2O$ | 46 ± NA | 1, S1 | [7] |
| 65% $H_2O$ - 35% PEG400 | 47.9 ± 0.3 | 1, S1, S2, S3 | [4] |
| 60% $H_2O$ - 40% PEG400 | 46.3 ± 0.3 | 1, S1, S2, S3 | [4] |
| 55% $H_2O$ - 45% PEG400 | 41.7 ± 0.3 | 1, S1, S2, S3 | [4] |
| 50% $H_2O$ - 50% PEG400 | 38.5 ± 0.3 | 1, S1, S2, S3 | [4] |
| 45% Dioxane - 55% $H_2O$ | 37 ± NA | 1, S1 | [7] |
| 40% $H_2O$ - 60% PEG400 | 33.1 ± 0.2 | 1, S1, S2, S3 | [4] |
| 30% $H_2O$ - 70% PEG400 | 27.4 ± 0.2 | 1, S1, S2, S3 | [4] |
| 20% $H_2O$ - 80% PEG400 | 23.1 ± 0.2 | 1, S1, S2, S3 | [4] |
| EtOH | 19 ± NA | 1, S1, S2, S3 | [14, 15] |
| 10% $H_2O$ - 90% PEG400 | 17.6 ± 0.1 | 1, S1, S2, S3 | [4] |
| ButOH | 16 ± NA | 1, S1 | [5] |
| 100% PEG400 | 14.7 ± 0.1 | 1, S1, S2, S3 | [4] |
| 100% hexan-1-ol | 10.3 ± N/A | 1, S1 | [5] |
| 12% anisol - 88% hexan-1-ol | 9.0 ± N/A | 1, S1 | [5] |
| 23% anisol - 77% hexan-1-ol | 7.9 ± N/A | 1, S1 | [5] |
| 34% anisol - 66% hexan-1-ol | 7.1 ± N/A | 1, S1 | [5] |
| 45% anisol - 55% hexan-1-ol | 6.3 ± N/A | 1, S1 | [5] |
| 55% anisol - 45% hexan-1-ol | 5.7 ± N/A | 1, S1 | [5] |
| 64% anisol - 36% hexan-1-ol | 5.3 ± N/A | 1, S1 | [5] |
| 74% anisol - 26% hexan-1-ol | 4.9 ± N/A | 1, S1 | [5] |
| 100% anisol | 4.5 ± N/A | 1, S1 | [5] |
| Mineral oil | 2.4 ± 0.04 | 1, S1, S2, S3 | [6] |

**Table S2:** Summary of the parameters of the calibration fits and interpolation functions of this work. All fits were performed by applying a $10^3$ weight factor to the first and last points of each calibration dataset to constrain the fit to capture both extremes of the permittivity interval. The relation $\frac{\phi(\lambda_f-\lambda_0)}{2\pi}+\lambda_0=\lambda$ was used to establish the link between the phase $\phi$ and the maximum emission wavelength $\lambda$ for the fit with Eq. 1 of the data presented in Fig. 1d and S1f.

| Symbol | Parameter | Value | Unit | Figures | Reference |
|---|---|---|---|---|---|
| $\alpha$ | 1st fitting parameter of Eq. S14 | $-3.74\times10^{5}$ | $cm^{-1}$ | 1c | This work |
| $\beta$ | 2nd fitting parameter of Eq. S14 | $9.74\times10^{-1}$ | | 1c | This work |
| $\gamma$ | 3rd fitting parameter of Eq. S14 | $2.16\times10^{6}$ | $cm^{-1}$ | 1c | This work |
| $\alpha$ | 1st fitting parameter of Eq. S14 | $-3.73\times10^{5}$ | $cm^{-1}$ | 1d | This work |
| $\beta$ | 2nd fitting parameter of Eq. S14 | $9.87\times10^{-1}$ | | 1d | This work |
| $\gamma$ | 3rd fitting parameter of Eq. S14 | $2.06\times10^{6}$ | $cm^{-1}$ | 1d | This work |
| $\alpha$ | 1st fitting parameter of Eq. S14 | $-7.31\times10^{3}$ | $cm^{-1}$ | S1e | This work |
| $\beta$ | 2nd fitting parameter of Eq. S14 | $8.45\times10^{-1}$ | | S1e | This work |
| $\gamma$ | 3rd fitting parameter of Eq. S14 | $2.50\times10^{4}$ | $cm^{-1}$ | S1e | This work |
| $\alpha$ | 1st fitting parameter of Eq. S14 | $-6.63\times10^{3}$ | $cm^{-1}$ | S1f | This work |
| $\beta$ | 2nd fitting parameter of Eq. S14 | $8.98\times10^{-1}$ | | S1f | This work |
| $\gamma$ | 3rd fitting parameter of Eq. S14 | $2.42\times10^{4}$ | $cm^{-1}$ | S1f | This work |
| $\mu_E$ | Excited state dipole moment | 20.9 | Debye | 1c-d | [3] |
| $\mu_G$ | Ground state dipole moment | 5.5 | Debye | 1c-d | [3] |
| $n_{H_2O}$ | Water refractive index | 1.335 | | 1c-d | [3] |

| Symbol | Parameter | Value | Unit | Figures | Reference |
|---|---|---|---|---|---|
| $V_{cav}$ | Volume of the dye virtual cavity | 0.2 | $nm^3$ | 1c-d | This work |
| $(\nu_f)_v$ | Emission wavenumber of ACDAN in vacuum | $2.44\times10^4$ | $cm^{-1}$ | 1c-d | This work |

**Table S3:** Comparing the static permittivity values reported in the literature for different solvents with the prediction of a simple law of mixture using the celebrated Maxwell-Garnett formula $\varepsilon_{eff} = \varepsilon_{H_2O}(2\phi_{EtOH}(\varepsilon_{EtOH} - \varepsilon_{H_2O}) + \varepsilon_{EtOH} + 2\varepsilon_{H_2O})/(2\varepsilon_{H_2O} + \varepsilon_{EtOH} - \phi_{EtOH}(\varepsilon_{H_2O} - \varepsilon_{EtOH}))$ where $\varepsilon_{eff}$ and $\phi_{EtOH}$ are the effective permittivity and the ethanol fraction in the mixture respectively. Note that the numerical value of ethanol permittivity used to compute the data in the 3rd column was set to $\varepsilon_{EtOH} = \varepsilon_{EtOH}(f \approx 1GHz) \approx 18.5\varepsilon_0$ (averaged from data available in refs. [14, 15]) where 1 GHz is approximately the inverse fluorescence lifetime of the ACDAN molecule. The data in the four rightmost columns associated with the present work were computed from the emission spectrum of ACDAN solvated in the relevant mixtures (1st column), acquired and processed via the referenced equipment and analytical approach; the indexes “Mic” and “Fluo” respectively relate to data acquisition performed with a microscope and spectrofluorimeter, while the mentions “DFT” and “Gauss” relate to the analytical approach (see Fig. 1b and Methods section) employed to process the measurements. Here and in the main text, all solvent values are vol%.

| Solvent | Literature | Law of mixtures | This work | | | |
|---|---|---|---|---|---|---|
| | $\varepsilon/\varepsilon_0$ from refs. [14, 15] | $\varepsilon/\varepsilon_0$ | $\varepsilon_{Mic}/\varepsilon_0$ (DFT) | $\varepsilon_{Fluo}/\varepsilon_0$ (DFT) | $\varepsilon_{Mic}/\varepsilon_0$ (Gauss) | $\varepsilon_{Fluo}/\varepsilon_0$ (Gauss) |
| 5% EtOH - 95% $H_2O$ | 73.2 | 74.3 | 74.9 ± 2.7 | 75.4 ± 2.9 | 74.8 ± 3.1 | 74.9 ± 3.4 |
| 10% EtOH - 90% $H_2O$ | 68-64 | 70.5 | 70.6 ± 2.6 | 71.3 ± 2.7 | 69.2 ± 2.8 | 71.7 ± 3.2 |
| 20% EtOH - 80% $H_2O$ | 57-55 | 63.0 | 61.1 ± 2.4 | 62.6 ± 2.4 | 55.9 ± 2.4 | 60.2 ± 2.7 |
| 30% EtOH - 70% $H_2O$ | 47-44 | 56.3 | 54.8 ± 2.2 | 53.9 ± 2.0 | 52.4 ± 2.2 | 50.5 ± 2.2 |
| 40% EtOH - 60% $H_2O$ | N/A | 49.8 | 47.4 ± 2.0 | 46.0 ± 1.7 | 46.2 ± 2.0 | 42.5 ± 1.9 |
| 50% EtOH - 50% $H_2O$ | 36 | 43.7 | 43.8 ± 1.9 | 41.9 ± 1.6 | 42.1 ± 1.8 | 37.4 ± 1.6 |
| 60% EtOH - 40% $H_2O$ | N/A | 38.0 | 38.4 ± 1.8 | 38.3 ± 1.5 | 37.4 ± 1.7 | 34.3 ± 1.5 |
| 70% EtOH - 30% $H_2O$ | 30 | 32.6 | 35.5 ± 1.7 | 34.4 ± 1.3 | 34.1 ± 1.5 | 31.5 ± 1.3 |
| 80% EtOH - 20% $H_2O$ | 29 | 27.4 | 29.5 ± 1.6 | 31.4 ± 1.2 | 29.6 ± 1.4 | 27.7 ± 1.2 |
| 90% EtOH - 10% $H_2O$ | 26 | 22.6 | 24.5 ± 1.4 | 27.1 ± 1.0 | 24.1 ± 1.2 | 22.5 ± 0.9 |
| 100% EtOH | 24 | 18.5 | 18.6 ± 1.2 | 22.95 ± 0.8 | 18.8 ± 1.0 | 18.7 ± 0.8 |

**Table S4:** Compositions of the PEG-rich and dextran-rich phases in the aqueous two-phase systems presented in Fig. 3a-b in the main text based on the phase diagram presented in ref. [16]. $w_p$ and $w_d$ are the respective total PEG and dextran weight fractions, $w_p^P$ and $w_d^P$ represent the PEG and dextran weight fractions in the PEG-rich phase, $w_p^D$ and $w_d^D$ are the PEG and dextran weight fractions in the dextran-rich phase, and $V_P$ and $V_D$ are the volume fractions occupied by the PEG-rich and the dextran-rich phases within the phase separated solution.

| Points, crosses | $w_p$(%) | $w_d$(%) | $w_p^P$(%) | $w_d^P$(%) | $w_p^D$(%) | $w_d^D$(%) | $V_P$(%) | $V_D$(%) |
|---|---|---|---|---|---|---|---|---|
| 1 | 3.0 | 4.8 | N.A. | N.A. | N.A. | N.A. | N.A. | N.A. |
| 2 | 3.2 | 5.1 | N.A. | N.A. | N.A. | N.A. | N.A. | N.A. |
| 3, 7 | 3.6 | 5.6 | 5.8 | 0.8 | 2.5 | 8.2 | 35 | 65 |
| 4, 8 | 3.8 | 6.0 | 6.3 | 0.6 | 2.0 | 9.9 | 42 | 58 |
| 5, 9 | 4.1 | 6.4 | 6.9 | 0.3 | 1.4 | 12.4 | 49 | 51 |
| 6, 10 | 4.3 | 6.8 | 7.4 | 0.2 | 1.1 | 13.8 | 51 | 49 |
